# Prediction of the intestinal resistome by a novel 3D-based method

**DOI:** 10.1101/196014

**Authors:** Etienne Ruppé, Amine Ghozlane, Julien Tap, Nicolas Pons, Anne-Sophie Alvarez, Nicolas Maziers, Trinidad Cuesta, Sara Hernando-Amado, Jose Luís Martínez, Teresa M. Coque, Fernando Baquero, Val F. Lanza, Luis Maiz, Tiphaine Goulenok, Victoire de Lastours, Nawal Amor, Bruno Fantin, Ingrid Wieder, Antoine Andremont, Willem van Schaik, Malbert Rogers, Xinglin Zhang, Rob J.L. Willems, Alexandre G. De Brevern, Jean-Michel Batto, Hervé Blottière, Pierre Léonard, Véronique Léjard, Aline Letur, Florence Levenez, Kevin Weiszer, Florence Haimet, Joël Doré, Sean P. Kennedy, S. Dusko Ehrlich

## Abstract

The intestinal microbiota is considered to be a major reservoir of antibiotic resistance determinants (ARDs) that could potentially be transferred to bacterial pathogens. Yet, this question remains hypothetical because of the difficulty to identify ARDs from intestinal bacteria. Here, we developed and validated a new annotation method (called pairwise comparative modelling, PCM) based on homology modelling in order to characterize the Human resistome. We were able to predict 6,095 ARDs in a 3.9 million protein catalogue from the Human intestinal microbiota. We found that predicted ARDs (pdARDs) were distantly related to known ARDs (mean amino-acid identity 29.8%). Among 3,651 pdARDs that were identified in metagenomic species, 3,489 (95.6%) were assumed to be located on the bacterial chromosome. Furthermore, genes associated with mobility were found in the neighbourhood of only 7.9% (482/6,095) of pdARDs. According to the composition of their resistome, we were able to cluster subjects from the MetaHIT cohort (n=663) into 6 “resistotypes”. Eventually, we found that the relative abundance of pdARDs was positively associated with gene richness, but not when subjects were exposed to antibiotics. Altogether, our results support that most ARDs in the intestinal microbiota should be considered as intrinsic genes of commensal microbiota with a low risk of transfer to bacterial pathogens.

## Introduction

Antimicrobial resistance is one of the major threats to health identified by the World Health Organization for the next decades^1^. The intestinal microbiota plays a pivotal role in this phenomenon as it harbours a vast diversity of bacterial species^2^, some of them possessing antibiotic resistance determinants (ARDs) that may enable their survival under antibiotic exposure^3^. Many of these ARDs did not evolve as “antibiotic resistance genes”, but provide a coincidental intrinsic resistance to antimicrobial drugs^4^. Previous studies have identified ARDs in the intestinal microbiota^5–7^ but the current bioinformatic tools for the annotation of ARDs, as well as in the case of other genes (based on sequence homology) are challenged by the important identity gap between known ARDs (mostly coming from culturable bacteria) and ARDs from the intestinal microbiota (mostly unculturable), that compromise the use of bioinformatic tools^8,9^. Thus, the real census of intestinal ARDs *i.e.* the intestinal resistome^10^ has not been comprehensively analysed to date.

While pathogenic bacteria have intrinsic, chromosomally-encoded ARDs and the capability of increasing resistance through mutation, they can also enrich their resistance capabilities through the acquisition of exogenous ARDs located on mobile genetic elements (MGEs) such as plasmids, transposons or phages. Thus, high-risk ARDs are commonly assumed to be highly mobile and widely transferable from various bacterial reservoirs to the pathogens^11^. Thereby the intestinal microbiota is assumed to be a potential reservoir of ARDs for bacterial pathogens, and even the origin of resistance determinants that have moved to such pathogens^12^. Still, despite the high selective pressure exerted on the intestinal microbiota by antibiotics that human have used for more than six decades, a very low number of transfer events from an intestinal commensal to a bacterial pathogen have been observed so far^13,14^. This could suggest that most ARDs from the intestinal microbiota are not located on MGEs, and thus less harmful in terms of public health since they would not substantially be fuelling pathogens for ARDs.

Previous studies have shown that individuals with no recent antibiotic exposure could be stratified according to the composition of their intestinal microbiota^15^. In agreement with the aforementioned hypothesis of a “mobile resistome”, stratifying subjects according to their intestinal resistome would not be feasible using the current bioinformatic tools. But if ARDs were to be mostly found located on the chromosome of their bacterial hosts, such stratification would be possible and would also open perspectives such as whether the microbiota of some subjects could resist the damage resulting from antibiotics exposure thanks to an inner important resistome when compared to subjects who would carry less ARDs. Indeed, whatever the route of administration or the purpose antibiotics may cause alterations in the composition of the intestinal microbiota and promotes the overgrowth of low-abundant, potentially pathogenic resistant bacteria over the susceptible ones^16,17^. Perturbations of the intestinal microbiota can be long-lasting, and could have consequences on health^18,19^. The scarce available data suggest that this effect is variable^16^, perhaps because of individual pharmacokinetics but also because of variations in the composition of the intestinal microbiota. Interestingly, some bacteria harbouring intrinsic ARDs might be involved in the protection of the intestinal microbiota towards antibiotics^20,21^, acting as “resilience” ARDs. Thereby, variable effects of antibiotics on the intestinal microbiota could possibly be explained by variations in the composition of the resistome.

Furthermore, the blossoming use of faecal material transplantation (FMT) or synthetic preparations of particular commensal bacteria (synthetic microbiota)^22,23^ calls for a careful description of the intestinal resistome and of the risk of ARDs transfers to bacterial pathogens.

To address those issues, we first searched for ARDs in the intestinal microbiota with a new functional annotation method and searched proteins associated with MGEs in their neighbourhood. We found that most ARDs we predicted (pdARDs) were distantly related to known ARDs and that they were not located on MGEs. Accordingly, we were able to cluster individuals with no recent exposure to antibiotics according to their resistome. We also found that the microbial gene richness was positively correlated to the abundance of the pdARDs, but this relationship was impacted by an exposure to antibiotics.

### Prediction of ARDs in the intestinal microbiota

To predict ARDs in the intestinal microbiota, we used a new method based on homology modelling (see methods) that we named PCM (for “pairwise comparative modelling”). PCM is a generic method using homology modelling to increase the specificity of functional prediction of proteins, especially when they are distantly related from proteins for which a function is known. The principle of PCM is to build structural models and assess their relevance using a specific training approach. PCM uses the list of sequences of reference proteins from a given family, the structures related to this family (they will be used as structural templates in the PDB format) and a series of negative references (Fig. 1A, Extended Data Fig. 1 and 2). The performances of PCM to predict ARDs were assessed with *in vitro* and *in silico* experiments. First, we performed *in vitro* experiments and synthesized 29 candidates for beta-lactamases further expressed in *Escherichia coli* (see methods). We detected a beta-lactamase activity in 20/29 (69.0%) of the tested candidates (Fig. 1B, see methods for the selection of candidates). The mean amino acid identity with known beta-lactamase of the 20 beta-lactamases successfully predicted by PCM and validated in vitro was 28.2% (range 19.4%-82.6%, Supplementary Table 1). Then, we used a functional metagenomics dataset from soils^24^, from which PCM could accurately identify 1,374 ARDs out of 1,423 hits (sensitivity 96.6%) with no false positive prediction (see methods). Eventually, we assessed the performances of PCM with incomplete proteins as inputs, and showed that PCM could correctly predict a function from proteins with >40% completeness (Extended Data Fig. 3). After the *in vitro* and *in silico* validation of the method, we used PCM to search for ARDs in the in a catalogue made of 3,871,657 proteins which was built from the sequencing of faecal samples of 396 human individuals (177 Danes and 219 Spanish) recruited in the MetaHIT project^25^ and for which metagenomic units (MGUs, referred to as clusters of co-abundant and co-occurring genes) have been determined^25^. In total, we predicted 6,095 ARDs (0.2% of the catalogue) from 20 ARD classes conferring resistance to nine major antibiotic families^26^: beta-lactams (classes A, B1-B2, B3, C and D beta-lactamases), aminoglycosides (AAC(2), AAC(3)-I, AAC(3)-II, AAC(6’), ANT, APH, RNA methylases), tetracyclines (Tet(M), Tet(X)), quinolones (Qnr), sulphonamides (Sul), trimethoprim (DfrA), fosfomycin (Fos) and glycopeptides (Van ligases) (Table 1 and Supplementary Table 1). With the same, extensively curated reference ARDs census as input, only 67 ARDs would have been predicted according to conventional search with a specific identity threshold (80%)^6,7^. Other specific tools Resfams^27^, ARG-ANNOT^28^ and Resfinder^29^ were able to predict 895, 54 and 50 ARDs, respectively (Fig. 1C). The mean identity shared between predicted and reference ARDs was 29.8%; it was significantly higher than candidates not predicted as ARDs (mean 23.0%, p<0.001, Figure 1D). Indeed, most of the pdARDs were distantly related to reference ARDs (Extended Data Fig. 4 and 5).

**Table 1:** Prediction of antibiotic resistance determinants (ARDs) from a 3.9M gene catalogue of the intestinal microbiota.^1^

| Family | Number of references | Number of candidates | Number of predictions of ARDs | Rate ARD predictions/candidates (%) |
| --- | --- | --- | --- | --- |
| Class A beta-lactamases | 682 | 402 | 267 | 66 |
| Class B1-B2 beta-lactamases | 150 | 554 | 134 | 24 |
| Class B3 beta-lactamases | 31 | 493 | 221 | 45 |
| Class C beta-lactamases | 56 | 373 | 76 | 20 |
| Class D beta-lactamases | 248 | 76 | 27 | 36 |
| AAC(2) | 5 | 15 | 3 | 20 |
| AAC(3)-I | 7 | 53 | 15 | 28 |
| AAC(3)-II | 12 | 81 | 81 | 100 |
| AAC(6) | 36 | 1191 | 312 | 26 |
| ANT | 29 | 158 | 67 | 42 |
| APH | 30 | 430 | 279 | 65 |
| RNA methylases | 17 | 4 | 2 | 50 |
| Tet(M) | 72 | 2824 | 1682 | 60 |
| Tet(X) | 12 | 42 | 9 | 21 |
| DfrA | 35 | 632 | 632 | 100 |
| Sul | 33 | 357 | 353 | 99 |
| Erm | 58 | 873 | 781 | 89 |
| Qnr | 66 | 272 | 219 | 81 |
| Fos | 34 | 84 | 62 | 74 |
| Van | 16 | 1163 | 873 | 75 |

**Figure 1:**
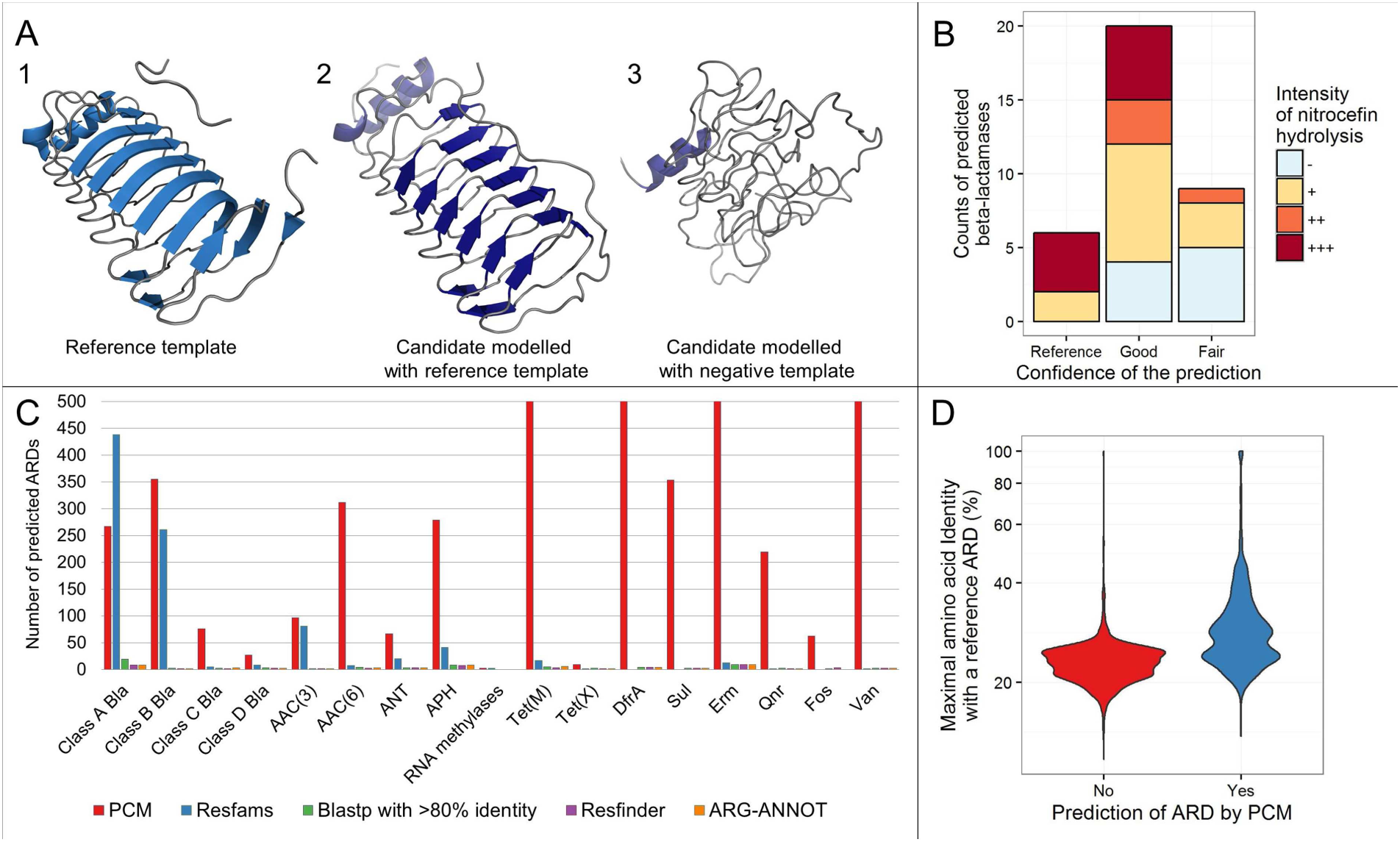
Illustration of the concept of “Pairwise Comparative Modelling” (PCM) with Qnr proteins (panel A). A1: Qnr protein structure (2W7Z) obtained from the PDB database. A2: A candidate protein (MC3.MG361.AS1.GP1.C32174.G14) for Qnr modelled with a reference Qnr structural template. This protein had 35.7% amino-acid identity with the closest Qnr protein (QnrB4). A3: The same candidate protein (MC3.MG361.AS1.GP1.C32174.G14) for Qnr this time modelled with a Qnr negative reference template. The candidate MC3.MG361.AS1.GP1.C32174.G14 was predicted to be a Qnr with 100% confidence by our model. Panel B: Bar-plot of the intensity of nitrocefin hydrolysis with respect to the degree of confidence of the prediction (“reference” meaning that the protein shares more ≥95% amino acid identity with a functionally proven beta-lactamase, “good” meaning a PCM score over 99% and a TmAlign Tm score ≥0.8, “fair” meaning a PCM score between 50% and 80%,). The intensity of the nitrocefin hydrolysis was determined in a semi-quantitative fashion, from duplicates experiments. Panel C: predictions of antibiotic resistance determinants (ARDs) from a 3.9M gene catalogue of the intestinal microbiota^1^ using PCM, Resfams^2^, Blastp, Resfinder^3^ and ARG-ANNOT.^4^ For Tet(M), DfrA, Erm and Vam, more than 500 predictions were obtained. Panel D: violin plot of the PCM prediction and maximal identity observed with a reference ARD. The y-axis is Log10 transformed. See Supplementary Table 1 for details about candidates sharing at least 40% identity with reference ARDs but were eventually not predicted as ARDs. AAC: aminoglycoside acetylase; ANT: aminoglycoside nucleotidyl transferase; APH: aminoglycoside phosphotransferase; DfrA: type A dihydrofolate reductase; Sul: dihydropteroate reductase; Erm: erythromycin ribosome methylase; Qnr: quinolone resistance; Fos: fosfomycin resistance (Fos); Van: D-Ala – D-Lac/Ser ligase (vancomycin resistance).

**Figure 2:**
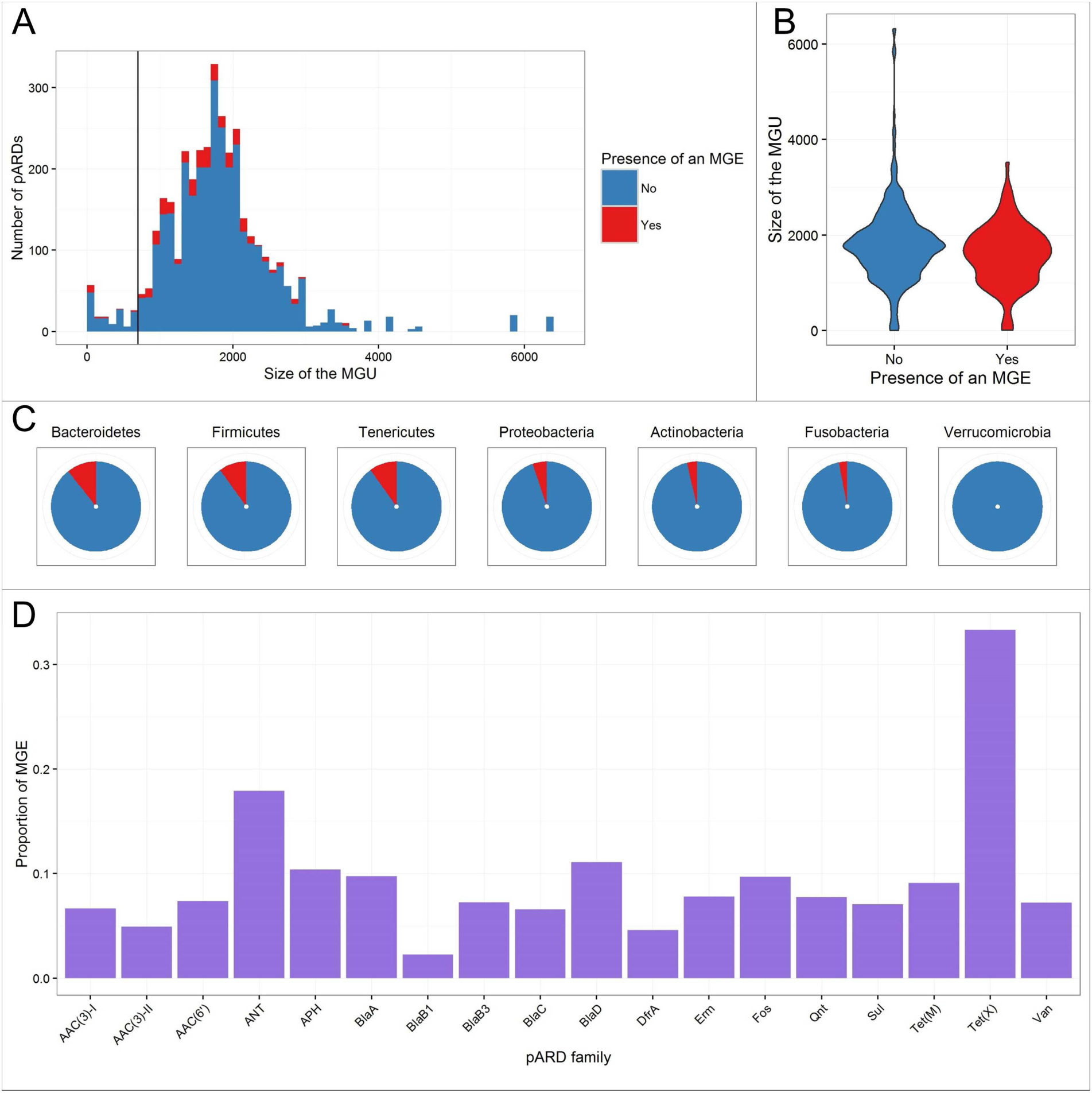
Mobile genetic elements (MGE) and predicted antibiotic resistance determinants (pdARDs). (A) Distribution of the sizes of the metagenomics unit (MGU) where an antibiotic resistance determinant was predicted with respect to the colocation of MGE-associated genes. The vertical line depicts the assumed gene size threshold above which MGUs are considered as partial chromosomes referred as metagenomic species (MGS)^1^. (B) Violin plots of the sizes of the metagenomics unit (MGU) where an antibiotic resistance determinant was predicted with respect to the colocation of MGE-associated genes. (C) Proportion of pdARDs co-locating with MGE-associated genes with respect to their phylum. (D) Proportion of pdARDs co-locating with MGE-associated genes according to the pdARD family. Of note, the AAC(2) and 16S RNA methylases only included 3 and 2 pdARDs, respectively and were accordingly not depicted in this panel.

**Figure 3:**
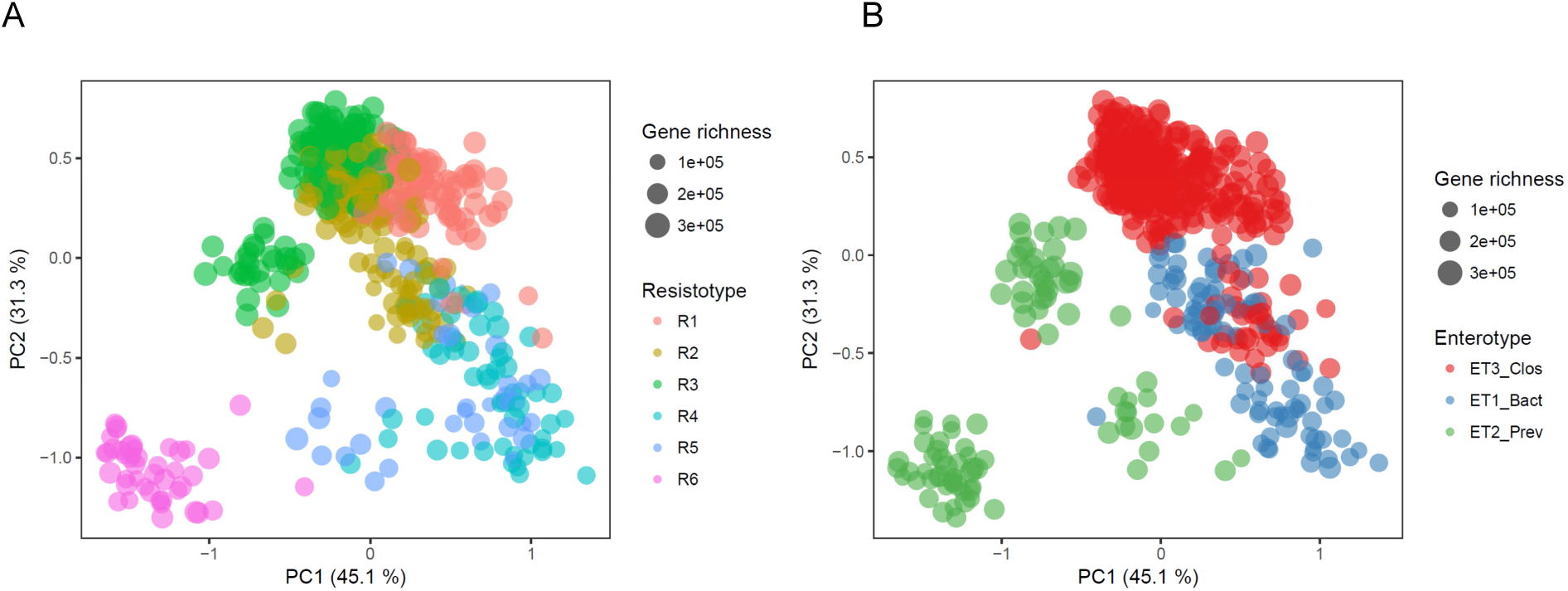
Principal component analysis of the 663 subjects of the MetaHIT cohort, with respect to their gene richness and resistotypes (A) or enterotypes (B).

**Figure 4:**
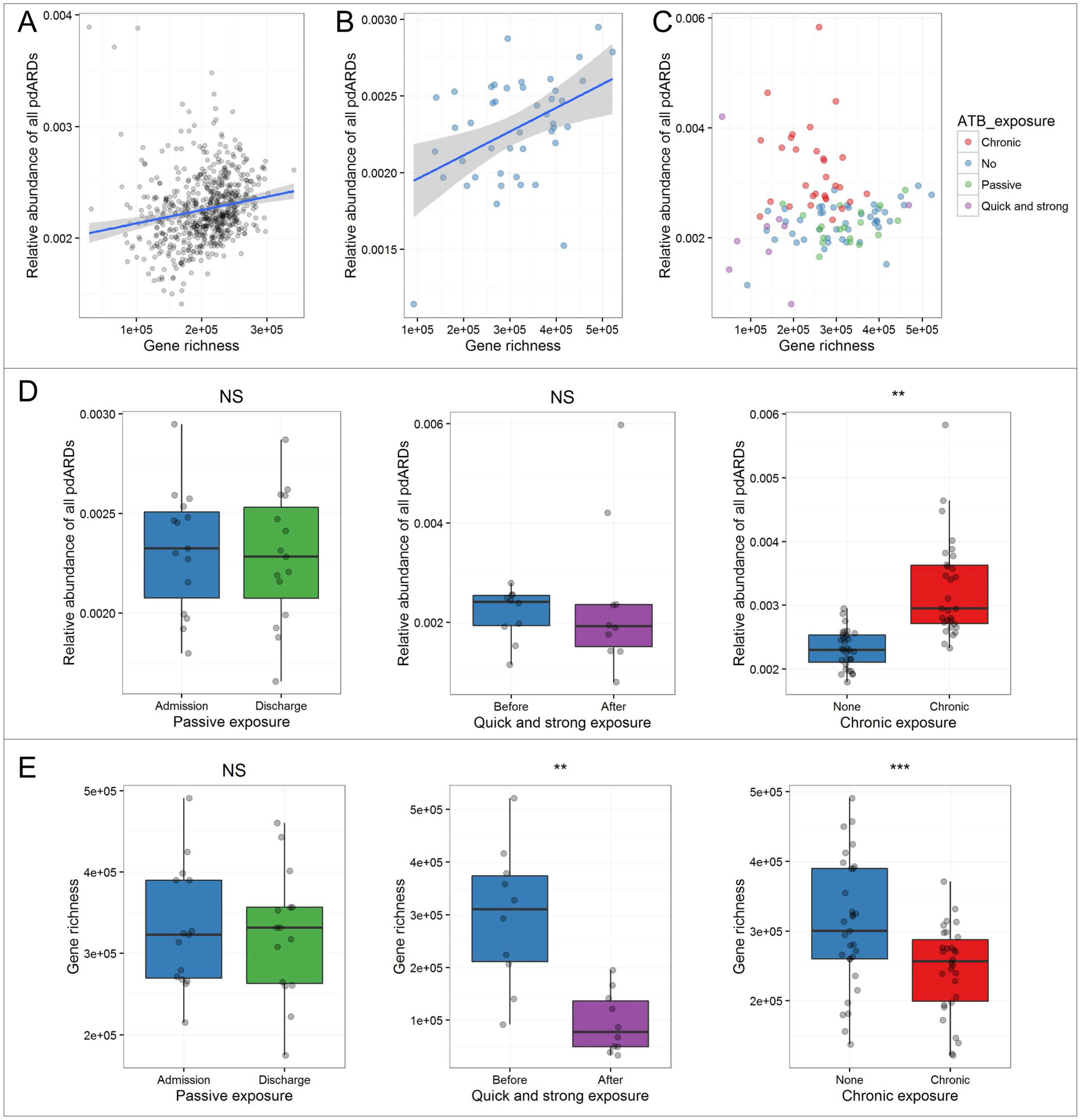
(A) Gene richness and relative abundance of predicted antibiotic resistance determinants (pdARDs) in the MetaHIT cohort (n=663). (B) Gene richness and relative abundance of pdARDs in our cohort of subjects with no recent antibiotic exposure (n=44). (C) Gene richness and relative abundance of pdARDs in our cohort of subjects with regards to their antibiotic exposure. (D) Boxplots superimposed by dot plots of the comparisons of the relative abundance of all pdARDs between the various groups differing by their exposure to antibiotics. (E) Comparisons between the gene richness between the various groups differing by their exposure to antibiotics. The unpaired Wilcoxon test was used. ***: p<0.001, ** p<0.01, *: p<0.05. The shaded grey area depicts the 95% confidence interval around the blue, linear regression line.

### Taxonomic distribution of ARDs

A host bacterial phylum could be attributed to 72.3% (4405/6095) pdARDs. The majority was identified as from Firmicutes (2962/4405, 72.3%) and from Bacteroidetes (858/4405, 19.5%) (Extended Data Fig. 6) with only 5.8% (225/4405) of pdARDs being from Proteobacteria. Of note, 7 pdARDs were predicted to be harboured by Archaea (*Methanobrevibacter* and *Methanoculleus* genera), putatively conferring resistance to macrolides, tetracyclines, aminoglycosides, sulphonamides and glycopeptides (Supplementary Table 1). Also, we predicted ARDs in genera of medical interest where no ARDs had been identified such as *Akkermansia*^30^ (10 pdARDs) and *Faecalibacterium*^31^ (44 pdARDs). Only 23 out of 6,095 (0.4%) had already been identified in families and genera that include human pathogens (Enterobacteriaceae, *Campylobacter*, *Enterococcus*, *Streptococcus* and *Acinetobacter*). Of note, PCM did not predict as ARD a total of 16 proteins sharing at least 40% identity with a reference ARD (Supplementary Table 2). pdARDs specifically distributed according to the taxa (Extended Data Fig. 7): Firmicutes and Proteobacteria were enriched with aminoglycosides-modifying enzymes (AMEs, spanning APH, ANT, and AACs) whereas Bacteroidetes were enriched in Sul and class A beta-lactamases. Interestingly, the tigecycline-degrading monooxygenase Tet(X) was frequently found in Bacteroidetes and Proteobacteria, the two phyla between which transfer of *tet(X)* gene has been reported^13,32^. Furthermore, we aimed to support these associations by *in vitro* experiments, and sequenced the metagenome of four human faecal samples before and after an overnight culture under conditions that favoured the growth of oxygen-tolerant bacteria such as Enterobacteriaceae and enterococci (see methods). We indeed observed a bloom of Proteobacteria (over Firmicutes and Bacteroidetes, Extended Data Fig. 8), in parallel to an increase of class C beta-lactamases, Fos and Tet(X), but also Van ligases (Extended Data Fig. 8).

### Location of the pdARDs and presence of neighbour mobile genetic elements

We investigated the mobility potential of the pdARDs at two levels. First, we assessed whether pdARDs were located on the chromosome or on a MGE. By using the 3.9M gene catalogue structuration into 7381 MGUs^25^, the 6095 pdARDS were assigned to their respective MGUs. Of note, MGUs which size exceeds 700 genes are assumed to be partial chromosomes and are referred as “metagenomic species” (MGS)^25^. Conversely, MGUs of smaller size can include MGEs such as plasmids, phages or transposable elements, but also incomplete chromosomal sequences. A total of 3,651 (59.9%) pdARDs could be mapped onto an MGU. The distribution of pdARDs according to the MGU gene content size is shown in Figure 2A. Most (95.6%, 3,489/3,651) pdARDs mapped onto MGS, strongly suggesting a chromosomal location. Then, we investigated whether genes associated with gene mobility such as transposases or conjugative elements were present in the genetic environment of the pdARDs. We first returned to the redundant catalogue (i.e. the non-dereplicated catalogue of genes)^25^ in order to obtain the homologues of the pdARDs and their respective contigs (n=16,955) containing a total of 908,888 genes (excluding the pdARDs). We searched for transposition and conjugative elements using ISFinder^33^ and Conjscan^34^ with sensitive parameters (see methods) and found that 7.9% (482/6,095) of pdARDs were co-located with genes putatively associated with gene mobility. Expectedly, we observed that the mean size of MGU that contained pdARDs co-locating with MGE were smaller than those who did not (Fig. 2B, p<0.001). Still, most MGE co-located with pdARDs were found in MGS (Fig. 2A). The phyla Bacteroidetes, Firmicutes and Tenericutes had the higher proportions of ARDs co-locating with MGEs (Fig. 2C). No ARD family was found to be enriched in MGE, with the exception of the Tet(X) family in which 3 out of 9 (33.3%) predictions (2 from *Bacteroides fragilis* and 1 from *E. coli*) were associated with transposases (Fig. 2D).

### Distribution of pdARDs in human hosts’ microbiota

From the MetaHIT cohort (663 subjects), we found that subjects carried pdARDs with a median relative abundance of 0.22% (range 0.14%-0.38%), with pdARDs from the Tet(M) family being the most abundant (0.07%) and those from class B3 beta-lactamases the least (median: 0.004%). The average number of unique pdARDs genes detected per metagenome was 1,377 (range 258-2,367). Most pdARDs were shared within subjects with only 106/6,095 (1.7%) with a single occurrence, and 987/6,095 (16.2%) found in at least 50% of individuals. All ARD families with the exception of RNA methylases and AAC(2’) families were found in more than 80% of individuals.

As pdARDs were found to be mostly chromosomal and not in the close neighbourhood of MGE-associated genes, we assessed whether subjects with no recent exposure to antibiotics could cluster according to their intestinal resistome as predicted by PCM. Based on the pdARDs family patterns, six clusters (that we named “resistotypes” by analogy with the enterotypes^15^) were detected using Dirichlet multinomial mixture models (Fig. 3A). The four most frequent resistotypes each represented around 20% of the cohort (the fifth and the sixth representing 8.7% and 7.5%, respectively). The three first resistotypes were characterized by an abundance enrichment of Van ligases (Extended Data Fig. 9), resistotype 1 was enriched in ANT, while resistotype 3 was driven by Tet(M) and class C beta-lactamases. Eventually, resistotype 4 was enriched with Tet(X) and class A beta-lactamases and resistotype 6 in class B1 beta-lactamases and Sul. We also observed that in the MetaHIT cohort, the relative abundance of pdARDs was positively correlated to the gene richness (Fig. 4A, Rho=0.3, p<0.001) and was associated with the enterotype classification (Fig. 3, chi square test, p<0.01). Resistotypes 1 and 3 had higher gene richness and were associated with the Clostridiales – driven enterotype. Resistotype 4 was more prevalent in enterotypes driven by Bacteroides (known to harbour Tet(X) and class A beta-lactamases) while resistotype 6 was very specific to the *Prevotella* enterotype (Fig. 3). Conversely, we did not find any link between resistotypes with body mass index, age or gender.

### Dynamics of the pdARDs under various exposures to antibiotics

Eventually, we investigated the variations of the abundances of pdARDs in subjects who experienced various exposures to antibiotics and healthcare environment. Three types of exposure were considered (see methods for details): passive (hospitalization in a French hospital without receiving antibiotics, n=17), chronic (Spanish cystic fibrosis patients frequently exposed to antibiotics, n=30) and quick and strong (selective digestive decontamination [SDD, made of oral colistin and tobramycin, and parenteral cefotaxime] at admission in intensive care units in Netherlands, n=10). We confirmed the positive relationship between relative abundance of pdARDs and gene richness among patients unexposed to antibiotics (Fig. 4B, Rho=0.37, p<0.05, see methods). When considering all the samples with regards to the recent exposure of the subjects, this relationship was not found anymore (Fig. 4C). Instead, we observed that the relative abundance of pdARDs was higher in subjects with a chronic exposure than in subjects with no recent exposure (Fig. 4D, p<0.01), while the gene richness was lower (Fig. 4E, p<0.001) In particular, subjects with chronic exposure carried more class B1-B2 beta-lactamases, AAC(6’), ANT, APH, Erm, DfrA while they had a lower abundance of Sul (Extended Data Fig. 10). At the phylum level, we observed a decrease of Bacteroidetes and Verrucomicrobia and an increase of Firmicutes and Actinobacteria in patients chronically exposed to antibiotics (Extended Data Fig. 11). A total of 74 MGS were found to be differentially abundant among subjects with or without chronic exposure to antibiotics (Supplementary Table 3).

This was different with subjects before and after they had SDD. We first observed a drastic loss of richness (Fig. 4E): from a mean of 295,919 genes to 95,286 (67.8 % reduction, Wilcoxon paired test, p=0.006). Meanwhile, the relative abundance of pdARDs did not significantly change (Fig. 4D, p=0.4). At the ARD family level, we observed that some bacterial families significantly decreased: class C beta-lactamases (commonly found in Enterobacteriaceae and Pseudomonadaceae which are the actual target of SDD), Fos, Tet(X), APH and ANT (Extended Data Fig. 12). We then analysed the MGS at the phylum level and found that Proteobacteria, Actinobacteria, Firmicutes and Fusobacteria significantly decreased after SDD (Extended Data Fig. 13). A total of 358 MGS were found in this cohort. Despite the small number of subjects (n=10), we found 133 MGS for which a significant variation was observed (a decrease after SDD in all cases but one, Supplementary Table 4). We tested whether a high abundance of pdARDs could be protective against the antibiotics used in SDD, but found not association: the relative abundance of pdARDs before SDD was not linked to the gene richness after SDD. Perhaps, as SDD uses a combination of antibiotics, bacteria that would be resistant to one antibiotic could be killed by one of the two others. Eventually, hospitalization without antibiotic therapy, that is, potential passive exposure to antibiotic resistant nosocomial organisms, did affect neither the gene richness nor the relative abundance of pdARDs, indicating selection as a key-factor in the microbiota incorporation of resistant microorganisms (Extended Data Fig. 4D and 4E).

## Discussion

Altogether, the results of this study support that the vast majority of ARDs from the intestinal microbiota could be considered as hosted by commensal bacteria, and that their transfer to an opportunistic pathogen is a rare event. We provide several findings to support this assumption: 1) we could assess the diversity of ARDs in the intestinal microbiota with a novel method that could predict functions for proteins which sequences were besides distant to known proteins, and confirmed that ARDs from the intestinal microbiota were distant from known ARDs from pathogens, 2) this method was validated by gene synthesis and application on an external, validation metagenomic dataset, 3) the majority of pdARDs showed no significant signature of genetic mobility, 4) this stability was illustrated by that we could stratify subjects into ‘resistotypes’, 5) we observed that gene richness, otherwise associated with a healthy status^19^, was positively correlated to the abundance of ARDs in subjects not exposed to antibiotics.

Our results indeed support that the threat to health by ARDs should be interpreted according to their genetic context, and strongly underlines the need of classifying ARDs accordingly to their risk for human health^11^. As it was previously demonstrated for soils^24^, ARDs tend to cluster by environment and the transfer of ARDs to pathogens occurs at a very low frequency. Our results suggest that the dominant intestinal microbiota (that is covered by metagenomic sequencing) is not a major reservoir that could fuel pathogens with ARDs, even though we acknowledge that such transfers did already occur^13,14^. Instead, the ARDs from our microbiota might not be a harm to public health, but they could furthermore act as potential protector against antibiotics as it was previously demonstrated for *Bacteroides* or *Prevotella* species producing beta-lactamases that protected the microbiota when a beta-lactam was given^20,21,35,36^. Our results call for further experiments to identify protective species with ARDs of low human risk concern^27^ that could potentially be included in synthetic microbiota as “bodyguards” for the bacteria of interest.

Our study has limitations, though. As mentioned earlier, metagenomic sequencing only covers the dominant bacteria, so that we could not cover the ARDs present in subdominant bacteria. Hence, we cannot rule out that the subdominant bacteria could be a reservoir of ARDs that could be transferred to pathogens. Besides, the method we used to identify distantly related proteins is based on homology modelling and takes advantage that proteins sharing the same function have more similar structures than amino acid sequences. While PCM showed excellent results on a soil dataset and on synthesized beta-lactamases, we nonetheless acknowledge it remains a prediction tool Indeed, two proteins sharing similar structures do not necessarily share the same functions and PCM shall yield false positive (as observed in the functional validation of synthesized beta-lactamases) so the number of ARDs in the intestinal microbiota may have been overestimated. Conversely, we are confident that PCM as it was built is a very sensitive method and that the true ARDs were identified. Indeed, PCM could identify functional beta-lactamases with an amino acid identity with known beta-lactamases below 20%. In all, we believe that PCM yields specific predictions but that in vitro experiments will remain necessary to confirm the functional annotation.

In all, we developed a novel method that could unveil the diversity of ARDs in the intestinal microbiota. Starting from this, we gathered evidence that the vast majority of the ARDs we predicted were, in majority, not mobile and that their abundance was correlated to the gene richness. Altogether, our results show that the ARDs from the intestinal microbiota could be considered as our “resilience allies”^37^ assuring the preservation of the healthy commensal microbiota under antibiotic exposure.

## Acknowledgements

The authors are deeply grateful to the GENOTOUL (Toulouse, France), GENOUEST (Rennes, France), ABIMS (Roscoff, France), MIGALE (Jouy-en-Josas) and TGCC-GENCI (Institut Curie) calculation clusters. The authors also warmly thank Bruno Perichon (Institut Pasteur, Paris, France) for providing ARD sequences from *Acinetobacter baumannii,* Patricia Siguier (CNRS, Toulouse, France) for helping the search of insertion sequences with ISfinder, Julien Guglielmini (Institut Pasteur, Paris, France) for his assistance in finding conjugative elements and Stevenn Volant (Institut Pasteur, Paris, France) for the design of the statistical model in SHAMAN.

## Methods

### Constitution of the databases of antibiotic resistance determinants

An ARD may be defined as a gene that decreases susceptibility to antibiotics when it is present or that increases susceptibility to antibiotics when it is lost^11^. This definition excluded intrinsic efflux pumps, target genes (in which mutations can confer resistance to some antibiotics) and genes involved in the regulation of antibiotic resistance genes. Amino acid sequences of functionally characterized ARDs from the major antibiotic families in human therapeutics (beta-lactam, aminoglycosides, tetracyclines, trimethoprim, sulfonamides, macrolides-lincosamides-synergistines, fluoroquinolones, fosfomycin and glycopeptides)^38^ were obtained from the following antibiotic resistance databases: Resfinder^29^, ARG-ANNOT^39^, the Lahey Clinic (http://www.lahey.org/studies/), RED-DB (http://www.fibim.unisi.it/REDDB/), Marilyn Roberts’ website for macrolides and tetracycline resistance (http://faculty.washington.edu/marilynr/) and from functional metagenomics studies^5,6,31^. Non-redundancy of the reference ARDs was assessed with CD-HIT v4.5.7^10^. The final database was manually curated. The cluster of orthologous genes (COG) of each member of the reference dataset was assigned from the v3 eggNOG database^40^. The mean size of the reference dataset sequences was determined. In total, we collected 1,651 non-redundant amino acid sequences spanning 20 ARDs families: Class A beta-lactamases (Blaa), class B1-B2 beta-lactamases (Blab1), class B3 beta-lactamases (Blab3), class C beta-lactamases (Blac), class D beta-lactamases (Blad), aminoglycosides acetyltransferases (AAC) AAC(2), AAC(3)-I, AAC(3)-II, and AAC(6), aminoglycosides nucleotidyltransferases (ANT), aminoglycosides phosphotransferases (APH), 16S rRNA methylases, Tet(M), Tet(X), type A dihydrofolate reductase (DfrA), dihydropteroate reductase (Sul), erythromycin ribosome methylase (Erm), quinolone resistance proteins (Qnr), fosfomycin resistance proteins (Fos), and D-Ala – D-Lac/Ser ligases (Van) (Table 1). The recently described plasmid-mediated colistin resistance *mcr-1* gene^41^ could not be included because of the lack of a reliable PDB template at the time of the study.

### Interrogation of the catalogue for ARDs

For each ARD family, we interrogated the 3,871,657 M proteins catalogue^16^ using three methods: (*i*) we built a hidden Markov model file for each ARD family and searched the catalogue with Hmmsearch (v3. 1)^42^, (*ii*) we performed a Blastp (v. 2. 2. 28+) search^43^ and (*iii*) a Smith-Waterman search (Ssearch v. 36. 3. 6)^44^ with an e-value threshold of 1E-5. Only the hits with a size ranging from 75% and 125% of the mean amino acid size of the ARD family were further considered. All candidates were assigned a COG/NOG from eggNOG v3. When candidates were found in different ARD families, the candidate was assigned the family including a reference for which it had the highest amino acid identity. We also searched for ARDs in the catalogue using conventional methods *i.e.* the same combination of Hmmsearch, Blastp and SSearch with both a minimal identity and a coverage over or equal to 80% of the candidate against a reference ARD.

### Negative references

For each ARD family, the COGs/NOGs attributed to candidates, that were not the COGs/NOGs of the reference dataset were further considered (Extended Data Fig. 2). We assumed that it reproduced the errors of functional assignment likely to be generated in sequence-only annotations. The hits from the Blast search against the eggNOG v3 database were added to the negative reference dataset. A manual curation step was performed in order to ensure that no references were eventually included in the negative references.

### Selection of structural templates

The list of protein structures that could be used as structural templates was downloaded (June 2014, and November 2014) from the PDB library (Protein DataBank^45^, http://www.rcsb.org/). Using the reference dataset and the negative references as found previously, Hmmer^46^, Blastp^43^ and SSearch^47^ were performed on the PDB database with default setting and e-values of 1E-5 and results merged into a non-redundant PDB list. Both lists (references and negative templates) were manually curated to ensure that no references were represented in the negative templates dataset, and reciprocally. If more than one PDB shared the same Uniprot number, we applied the following criteria: no ligand, completeness of the protein and high resolution in order in include a unique structure per Uniprot number.

### Pairwise comparative modelling

The concept of pairwise comparative modelling (PCM) is shown in Extended Data Fig. 1 and 2, and the framework is available at https://github.com/aghozlane/pcm. Each candidate was subjected to homology modelling with reference templates and negative templates, generating two 3D-structures for each candidate (Fig 1A). The main idea is that if the sequence is really associated to the reference fold, its model must be significantly different from the ones obtained with the negative structural template. Homology modelling was performed by PCM in seven main steps:

1. Three structural templates were identified by Blastp (among the lists produced as described above) that shared the highest homology with the candidate protein.
2. A multiple sequence alignment was performed between the candidate and the three templates sequences using Clustalo^48^.
3. A prediction of the secondary structure was performed using psipred (v3.5)^49^. The residues predicted to fold in helix or in beta-sheet conformation with a level of confidence higher or equal to 7 were considered to constrain the model that will be produced.
4. A comparative modelling was performed with the MODELLER programming interface^50^. MODELLER automatically calculates a model by satisfaction of spatial restraints such as atomic distance and dihedral angles in the target sequence, extracted from its alignment with the template structures. Stereo-chemical restraints for lasting residues are obtained from the CHARMM-22 molecular force field and statistical preferences obtained from a representative set of known protein structures.
5. The best model out of a hundred produced by MODELLER (based on the Dope score) was considered for structure assessment analysis using ProQ^51^ and Prosa-web^52^. The Dope score (Modeller), z-score (Prosa), MaxSub and LG score (ProQ) are statistical potential variables used to predict the model quality. Theses scores are trained on the PDB and they estimate the energetic favourability of the conformation of each residue in the model.
6. The best model was aligned with the reference set of structures using TM-align^16^ and MAMMOTH^53^. The RMSD (TM-align), z-score (MAMMOTH), TM-score (MAMMOTH, TM-align) estimates the degree of superposition of the residue between two structures. The differences (delta) between the scores determined from each modelling path (with the reference set or the negative set) were calculated and used for the PCM machine learning program (see below). For one given candidate, the PCM whole process took an average of 8 CPU-hours (30 minutes on 16 CPUs).

### Statistical analysis

To discriminate reference proteins from negative references, we used model quality predictors and alignment scores (inferred from the semi-automatic pipeline described in the section above) and developed a custom pipeline in R (R Core Team, 2013, http://www.R-project.org) to perform the classification. The LASSO penalized logistic regression^54^ implemented in LIBLINEAR^55^ was used to compute the classifier. Ten-fold stratified cross validation (re-sampled 100 times to obtain more stable accuracy estimates) was used to partition the data into a training set and a test set. The LASSO hyper-parameter was optimized for each model in a nested 5-fold cross-validation on the training dataset using the area under curve (AUC) as model selection criterion. From the 100 times re-sampled ten-fold cross validation, receiver operating characteristic (ROC) analysis was used to evaluate the model performances using the median AUC. Coefficients extracted for each modelling or alignment score were also evaluated for their stability throughout the computed models. The PCM score was the ratio (expressed as a percentage) between the numbers of time a candidate was classified as a reference and the number of bootstraps. Predicted ARDs were candidates with a PCM score over or equal to 50% and a TM score given by TM-align over or equal to 0. 5^16^.

### Validation of the method with a functional metagenomic dataset

In order to assess the performances of PCM, we took advantage of the publication by Forsberg *et al*. in 2014 where the ARD content of different North American soils was analysed using functional metagenomics^24^. The screening of the clones was performed on aztreonam, chloramphenicol, ciprofloxacin, colistin, cefepime, cefotaxime, cefoxitin, D-cycloserine, ceftazidime, gentamicin, meropenem, penicillin, piperacillin, piperacillin-tazobactam, tetracycline, tigecycline, trimethoprim and trimethoprim-sulfamethoxazole (cotrimoxazole). For the present study, we collected the nucleotide sequences of the inserts deposited on Genbank (KJ691878–KJ696532). The sequence translation of the open reading frames was performed by Prodigal (using default parameters)^56^. A total of 4,654 sequences of inserts were collected, in which 12,904 amino acid sequences were predicted. We then searched for ARDs belonging to the relevant ARD families according to the antibiotics used for the screening of the clones: beta-lactamases (all classes), APH, ANT, AAC(2), AAC(3)-I, AAC(3)-II, AAC(6), RNA methylases, Tet(M), Tet(X), Qnr, Sul and DfrA, using the supplementary table 2 of the paper. Inserts with no ARDs were removed (n=269). Also, inserts selected on cycloserine (n=868) and chloramphenicol (n=129) were not considered here. Fourteen inserts which contained more than one ARD that could be predicted to confer resistance to the antibiotic used for the screening (e. g.; two beta-lactamases) were not considered in this analysis. Eventually, 1,658 inserts containing no identified ARDs or ARDs that did not confer resistance to the antibiotic used for selection were discarded and 294 containing efflux pumps (not considered in this study). Thus, the validation set was made of 1,423 inserts (containing a resistance gene) for a total of 3,778 genes. Class B1-B2 and class B3 beta-lactamases were modelled separately then doublets were removed and results were shown as class B beta-lactamases as the precise class annotation was not provided in the dataset.

In total, 1,390 unique hits were found during the initial screen of PCM, of which 1,374 were predicted as ARDs (Supplementary Table 5). Among the 33 ARDs not included for PCM, 12 were not considered because they were undersized and 10 because they were oversized. No hits for AAC(2), ANT, Qnr or Sul were found. The mean identity shared with reference ARDs was 37.6% (range 18.8-94.5). Overall, the sensitivity and negative predictive value were 96.6% and 98.0%, respectively. In comparison, only 8 ARDs would have been identified by a conventional method (combination of Hmmsearch, Blastp and SSearch with both a minimal identity with a reference ARD and coverage over or equal to 80%). Conversely, Resfams^27^ that was specifically designed to identify ARDs from functional metagenomic datasets showed similar sensitivity to PCM with 1,346 correct predictions of ARDs (94.6% sensitivity).

### Validation of the method for incomplete genes

The 3.9M gene catalogue harbours 41.4% of genes that are predicted to be incomplete either on the 5', the 3' or both extremities^25^. As the size parameter is crucial for homology modelling, we tested to which extent the prediction of incomplete ARDs by PCM could remain correct. We selected 12 reference class A beta-lactamases (BlaZ, CblA-1, CepA-29, CfxA2, CfxA6, CTX-M-8, KPC-10, OXY-1, PER-1, SHV-100, TEM-101 and VEB-1) for that we iteratively removed 5% of the amino acid sequence at both edges in order to obtain 16 bi-directionally trimmed candidates (from 100% to 25%) per reference ARD. A total of 192 PCM were performed: we observed that the 12 references were correctly predicted as ARDs when the trimming proportion degree was over 40% (*i.e.* 30% trim from each extremity, Supplementary Data Fig. 3). Thus, we are confident that with the 75% size threshold we chose (25% maximal cut from one edge), no misclassification due to an incompleteness gene could be expected.

### Gene synthesis

We selected 35 predicted beta-lactamases (14 from class A, 8 from class B1-B2, 7 from class B3, 4 from class C and 2 from class D) for gene synthesis and sub-cloning into *E. coli* to test the beta-lactamase function. We used the plasmid vector pET-26b+ (embedding a kanamycin resistance - conferring gene). The selection of the predicted beta-lactamases for synthesis was performed as follows:

- Reference beta-lactamases (n=6): ≥95% amino acid identity and ≥80% coverage with known ARDs.
- Beta-lactamases with the highest degree of confidence for the prediction (n=16): PCM score >99%, Tm score TmAlign>0.9 and <70% amino acid identity with a reference beta-lactamase.
- Beta-lactamases with the lowest degree of confidence for the prediction (n=8): PCM score <80%, Tm score TmAlign<0.8.
- Beta-lactamases randomly selected (n=5)

The *E. coli* transformed with the plasmids embedding the predicted beta-lactamases were then tested for their activity in using a broth culture containing nitrocefin as chromogenic indicator of beta-lactamase activity. The degree of hydrolysis was semi-quantified according to the intensity of the coloration: negative (pink), + (orange), ++ (dark orange) and +++ (yellow). Of note, we chose to synthesize beta-lactamases to test the hypothesis that secreted ARDs could detoxify antibiotics in the gut (see results). As the candidates ranged into two main families that are structurally distinct (class A, C and D being serine beta-lactamases and class B being metallo-enzymes), we assume that the observations from this experiment could apply to the other non-beta-lactamases families.

### Searching for signatures of mobile genetic elements nearby the predictions of ARDs

Mobile genetic elements (MGE) - associated proteins were searched in the genetic environment of the pdARDs. First, homologues of insertion sequences were searched in the redundant gene catalog, contigs where an ARD was predicted (n=908,888) using ISfinder^33^ and blastp (query size threshold 150 amino acids, e-value 1E^-30^, identity threshold 40%) against the ISfinder database. Furthermore, conjugative elements were searched among the same gene set (n=908,888) with Conjscan^34^, using the default parameters and the filters recommended by the authors (best e-value<0.001 and sequence coverage of at least 50%). A pdARD was considered to be linked to a conjugative element when at least two distinct elements and/or one mob element were found in the contig(s) of its homologues (irrespective of the genetic distance between the pdARD gene and the MGE-associated gene).

### Distribution of the ARDs in the MetaHIT cohort (n=663 subjects)

pdARDs profiles were obtained from the abundance matrix of the 3.9M genes as described in Nielsen *et al*^25^. The “reads per kilobase per million mapped reads” (RPKM) method was used to normalize the mapping counts. After summing the relative abundances of pdARDs genes belonging to the same family, Dirichlet multinomial mixture models were used to find ARDs clusters (*i.e.* resistotypes) using the Dirichlet Mulitnomial R package. The same methods was applied to detect gut microbiota clusters (*i.e.* enterotypes)^57^. Laplace criterion was used to define optimal number of clusters as described on oral and faecal microbial dataset^58^. By analogy with the term enterotype, we chose to name as “resistotype” a cluster of subjects based on their similarity of their faecal relative abundance of pdARDs. Chi-squared test was used to assess the associations between resistotypes and enterotypes. Rarefaction analysis at one million read was done to determine the gene richness per samples. RLQ analysis^59^ was conducted to assess the associations between the relative abundances of pdARDs, their characteristics (family, size of the cluster of associated genes [CAG]) and those of subjects (enterotypes, resistotypes, gender, body mass index [BMI], age). Of note, we excluded the patients suffering from inflammatory bowel disorders from this analysis.

### Constitution of cohorts of patients with various antibiotic exposures

We included three cohorts of patients with various exposures to antibiotics:

- Passive exposure: a total of 32 patients with no exposure to antibiotics or hospitalisation during the preceding 3 months and admitted to the medicine ward of the Beaujon University Teaching Hospital (Clichy, France) were included and provided a faecal sample. Among them, 17 could provide a stool at discharge. Two received some antibiotics during their stay and their samples at discharge were not considered for the analysis. In total, 15 patients could provide a stool sample soon after admission (T0) and at discharge (T1). The mean duration of passive exposure (mean time between T0 and T1 samples) was 10. 7 days. We also could include 13 subjects who could provide a sample at admission and who had not been exposed to antibiotics in the three months preceding their visit.
- Chronic exposure: 30 cystic fibrosis (CF) - suffering patients were enrolled at the Cystic Fibrosis Unit of the Ramon y Cajal Hospital in Madrid, mostly non-hospitalized. Cystic fibrosis is a genetic disease that leads to an impairment of the lung function through an uncontrolled production of mucus. The consequence is chronic bacterial colonization, resulting in deleterious reactive fibrosis of the lung. Bacterial load is controlled by chronic exposure to antibiotics (home-therapy, mostly oral and inhaled in our cohort), which has resulted in significant life prolongation, and almost absence of need of hospital care. One faecal sample was collected at the occasion of a consultation.
- Quick and strong: selective digestive decontamination (SDD) consists in administering an association of topical and parenteral antibiotics and antifungal agents to a patient at admission in order to eliminate potential bacterial and fungal pathogens. SDD has been showed to significantly reduce mortality in the intensive care unit (ICU)^60^ and is now part of standard care in Netherlands. Still, the selection for antibiotic-resistant pathogens by SDD remains a concern, especially resistance to colistin^61^ which is a last-resort antibiotic in infections caused by extensively-resistant Gram-negative bacilli. To assess the effect of SDD on the intestinal microbiota, we analysed the faecal samples from 13 patients admitted to the ICU of the University Medical Center of Utrecht (UMCU, Netherlands). The samples were collected at admission (T0, first sample passed after admission) and after SDD (T1). Among the 13 patients for whom a faecal sample could be obtained at T0, 10 could provide a faecal sample at T1. SDD consisted in of 4 days of intravenous cefotaxime and topical application of tobramycin, colistin, and amphotericin B. Besides, a subset of samples (n=4) from this cohort were cultured in a brain-heart infusion broth overnight in ambient atmosphere at 37°C.

### Metagenomic sequencing and mapping

Total faecal DNA was extracted^51,52^ and sequenced using SOLiD 5500 wildfire (Life Technologies) resulting in a mean of 68.5M sequences of 35-base-long single-end reads. High-quality reads were generated with quality score cut-off >20. Reads with a positive match with human, plant, cow or SOLiD adapter sequences were removed.

Filtered high-quality reads were mapped to the MetaHIT 3.9M gene catalog ^25^ using the METEOR software^64^. The read alignments were performed in colorspace with Bowtie software (version 1.1.0) ^65^ with options: -v 3 (maximum number of mismatch) and -k 10000 (maximum number of alignment per reads). The gene abundance profiling table was generated using a two-step procedure. First, the unique mapped reads (reads mapping to a unique gene from the catalogue) were attributed to the corresponding genes. Then the shared reads (mapping different genes of the catalogue) were attributed according to the ratio of their unique mapping counts, as following: as a read can map on different genes of the catalogue, the abundance of a gene *G(A_g_)* depends on the abundance of uniquely mapped reads (*A_u_*), *i.e.* reads that map only to the gene *G*, and on the abundance of *N* shared reads (*A_s_*) that aligned with *M* genes in addition to the gene G:

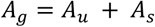

Where

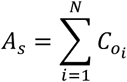

For each shared read, the gain of abundance corresponds to a coefficient *C_0_* that takes in account the total number of uniquely mapped reads on the *M* genes:

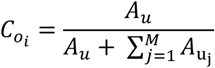

For instance, if a gene G is mapped by 10 reads that only map to it (unique reads), but also with 1 read that also align on a gene M that was mapped by 5 unique reads, then:

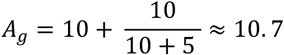

To decrease technical biases due to different sequencing depth, samples with at least 5M mapped reads were downsized to 5M mapped reads (random sampling of 7M mapped reads without replacement) using R package momr^19^. The abundance of each gene in a sample was then normalized by dividing the number of reads that mapped to the gene *(A*_*g*_) by the gene nucleotide length and by the total number of reads from the sample. The resulting set of gene abundances, termed a “microbial gene profile”, was used to estimate the abundance of metagenomic species (MGS). ^25^

#### Gene richness analysis

Microbial gene richness was calculated by counting the number of genes that have been mapped at least once in a given sample. Of note, a high microbial gene richness has been associated to be a marker of a healthy gut microbiota, even if a specific threshold remains to be set^19^. Gene richness was calculated using R package momr for samples where 5M or more reads had mapped against the 3. 9M catalogue.

#### MetaGenomic Species (MGS)

MGS are co-abundance gene groups with more than 700 genes, and can thus be assumed as part or complete bacterial species genes contents. 741 MGS were delineated from 396 human gut microbiome samples^25^. In this study, the relative abundance of MGS were determined as the median abundance of 90% of the genes composing each cluster. MGS taxonomical annotation was updated by sequence similarity using NCBI BLASTN, when more than 50% of the genes matched the same reference of NCBI database (December 2014 version) at a threshold of 95% of identity and 90% of gene length coverage to get the species annotation.^25^

#### Statistical analysis for the distribution of pdARDs and MGS between groups

Statistical analysis for the differential abundances of ARDs and MGS in the patient cohorts were performed using the application SHAMAN^66^ (http://shaman.c3bi.pasteur.fr/, data are available at https://github.com/aghozlane/evotar). The relationship between richness and the abundance of ARDs was assessed by the Spearman correlation test. The statistical threshold was set at a p-value of 0.05. The p-values were represented by a single * when 0.01<p<0.05, ** when 0.001<p<0.01 and *** when p<0. 001.

